# Structural basis for EPC1-mediated recruitment of MBTD1 into the NuA4/TIP60 acetyltransferase complex

**DOI:** 10.1101/746180

**Authors:** Heng Zhang, Maëva Devoucoux, Xiaosheng Song, Li Li, Gamze Ayaz, Harry Cheng, Wolfram Tempel, Cheng Dong, Peter Loppnau, Jacques Côté, Jinrong Min

## Abstract

MBTD1, a H4K20me reader, has recently been identified as a component of the NuA4/TIP60 acetyltransferase complex, regulating gene expression and DNA repair. NuA4/TIP60 inhibits 53BP1 binding to chromatin through recognition of the H4K20me mark by MBTD1 and acetylation of H2AK15, blocking the ubiquitination mark required for 53BP1 localization at DNA breaks. The NuA4/TIP60 non-catalytic subunit EPC1 enlists MBTD1 into the complex, but the detailed molecular mechanism remains incompletely explored. Here, we present the crystal structure of the MBTD1-EPC1 complex, revealing a hydrophobic C-terminal fragment of EPC1 engaging the MBT repeats of MBTD1 in a site distinct from the H4K20me binding site. Different cellular assays validate the physiological significance of the key residues involved in the MBTD1-EPC1 interaction. Our study provides a structural framework for understanding the mechanism by which MBTD1 recruits the NuA4/TIP60 acetyltransferase complex to influence transcription and DNA repair pathway choice.

## Introduction

Coordination of posttranslational modifications (PTMs) represents an important signalling mechanism to control often complex biological processes such as DNA repair (Chin, 2017; Dantuma and van Attikum, 2016; Woodsmith et al., 2013; Woodsmith and Stelzl, 2014). DNA double strand break (DSB) is one of the most severe forms of genomic lesions whose improper repairs will trigger genomic instability and lead to malignant transformation (Hustedt and Durocher, 2016; Pommier et al., 2016; Wilson and Durocher, 2017). To properly repair these dangerous genetic lesions, mammalian cells have evolved complex yet highly orchestrated signalling cascades involving various PTMs such as ubiquitylation, methylation and acetylation for context-dependent signal propagation. The NuA4 (Nucleosome acetyltransferase of H4) /TIP60 acetyltransferase complex has been demonstrated to participate in the repair of DNA DSBs at multiple levels, underlining the pleiotropic contributions of acetylation as a PTM to the DNA DSB repair network (Dhar et al., 2017; Jacquet et al., 2016; Paquin and Howlett, 2018; Rossetto et al., 2010; Soria et al., 2012; Tang et al., 2013).

Two major pathways exist to repair DNA DSBs in mammalian cells, namely homologous recombination (HR) and non-homologous end joining (NHEJ) (Her and Bunting, 2018; Sunada et al., 2018). The choice between these two canonical DSB repair pathways often determines the cellular fate post DNA damage and can have significant implications for related human syndromes as well as therapeutic strategies (Chapman et al., 2012; Liu et al., 2016; Sung, 2018). The recruitment and retention of 53BP1 has been suggested to potentiate NHEJ-mediated repair of DSBs primarily by safeguarding DSB ends from DNA end resection, a key requirement to initiate HR-mediated repair (Gupta et al., 2014; Liu and Huang, 2016). While 53BP1 can be recruited to the DSB sites in part via binding to the H4K20me mark, RNF168 mediated mono-ubiquitination on H2AK13/15 is also key to the 53BP1 appearance at the break, leading to the DSB being repaired through the error-prone NHEJ pathway (Fradet-Turcotte et al., 2013; Zimmermann and de Lange, 2014). Another H4K20me reader protein, MBTD1 (MBT domain-containing protein 1), has recently been demonstrated to be a stable subunit of the NuA4/TIP60 acetyltransferase complex (Jacquet et al., 2016). Interestingly, MBTD1 affects the recruitment of the NuA4/TIP60 complex to a specific subset of genes to regulate their expressions (Jacquet et al., 2016). NuA4/TIP60-associated MBTD1 binding to the H4K20me mark at DSB sites competes with 53BP1 for association with the chromatin surrounding the lesion (Jacquet et al., 2016). The MBTD1-linked Tip60 (Tat interacting protein 60) enzyme catalyzes acetylation of not only the histone H4 tail, destabilizing 53BP1 interaction (Tang et al., 2013), but also H2AK15 to prevent its ubiquitylation by RNF168, and thereby further blocking the recruitment and retention of 53BP1 to the DSB sites (Jacquet et al., 2016). Collectively, MBTD1 via its direct interaction with the EPC1 (Enhancer of polycomb homolog 1) subunit of the NuA4/TIP60 acetyltransferase complex negatively regulates 53BP1’s bivalent associations with chromatin surrounding DSB sites through competition for binding to H4K20me and histone lysine acetylation (Jacquet et al., 2016). Therefore, this MBTD1-EPC1 interaction not only regulates certain gene expressions, but also enables the recruitment of the DNA end resection machinery to the DSB site and commits the cellular response to the error-free HR pathway.

MBTD1 consists of a N-terminal FCS zinc finger that binds to regulatory RNAs and four malignant brain tumor (MBT) domains that recognize H4K20me1/me2 mark (Eryilmaz et al., 2009; Lechtenberg et al., 2009). The first and second MBT domains are reported to accommodate transcription factor YY1 in vitro (Alfieri et al., 2013), whereas the fourth MBT domain harboring the semi-aromatic cage is responsible for histone mark readout (Eryilmaz et al., 2009). EPC1 contains EPcA, EpcB and EPcC domains at the N-terminus (Kee et al., 2007). The C-terminus of EPC1 was reported to interact with transcriptional repressor RFP (Tezel et al., 2002). The precise molecular mechanism by which MBTD1 associates with EPC1 remains elusive. Through structural analysis coupled with biophysical and functional analysis, we have elucidated the interaction mechanism between MBTD1 and the EPC1-containing NuA4/TIP60 acetyltransferase complex. Hence, these data will provide valuable information for the development of inhibitors for dissociating TIP60 acetyltransferase from the DSB sites, and thereby will enable the manipulation of DSB repair pathway choice that can have clinical implications.

## Results and Discussion

### Characterization of the key EPC1 region responsible for binding to MBTD1

MBTD1 has been recently revealed to associate with the NuA4/TIP60 acetyltransferase complex via its interaction with the C-terminus of EPC1, a non-catalytic subunit of the core complex (Jacquet et al., 2016). To investigate the detailed molecular mechanism behind the MBTD1-mediated recruitment of the NuA4/TIP60 acetyltransferase complex to chromatin, we first aimed to narrow down the key region on the C-terminus of EPC1 (**Figure 1A**). We performed *in vitro* GST pull-down experiments followed by western blot. Notably, GST-EPC1_644-799_ efficiently bound MBTD1, whereas GST-EPC1_729-799_ and GST-EPC1_673-776_ lacking a hydrophobic region did not (**Figures 1A, 1B and S1**), suggesting that this hydrophobic fragment (residues 644-672) of EPC1 is likely responsible for binding of MBTD1. Indeed, the GST-EPC1_644-672_ fragment was able to pull down MBTD1 (**Figure 1B**). Further Isothermal Titration Calorimetry (ITC) studies confirmed that EPC1_644-672_ directly interacts with MBTD1 with a dissociation constant (K_d_) of ∼67 nM (**Figure 1C**), indicating that the hydrophobic C-terminal fragment of EPC1 (residues 644-672) enlists MBTD1 into the dynamic NuA4/TIP60 acetyltransferase complex (**Figure 1A**).

**Figure 1.**
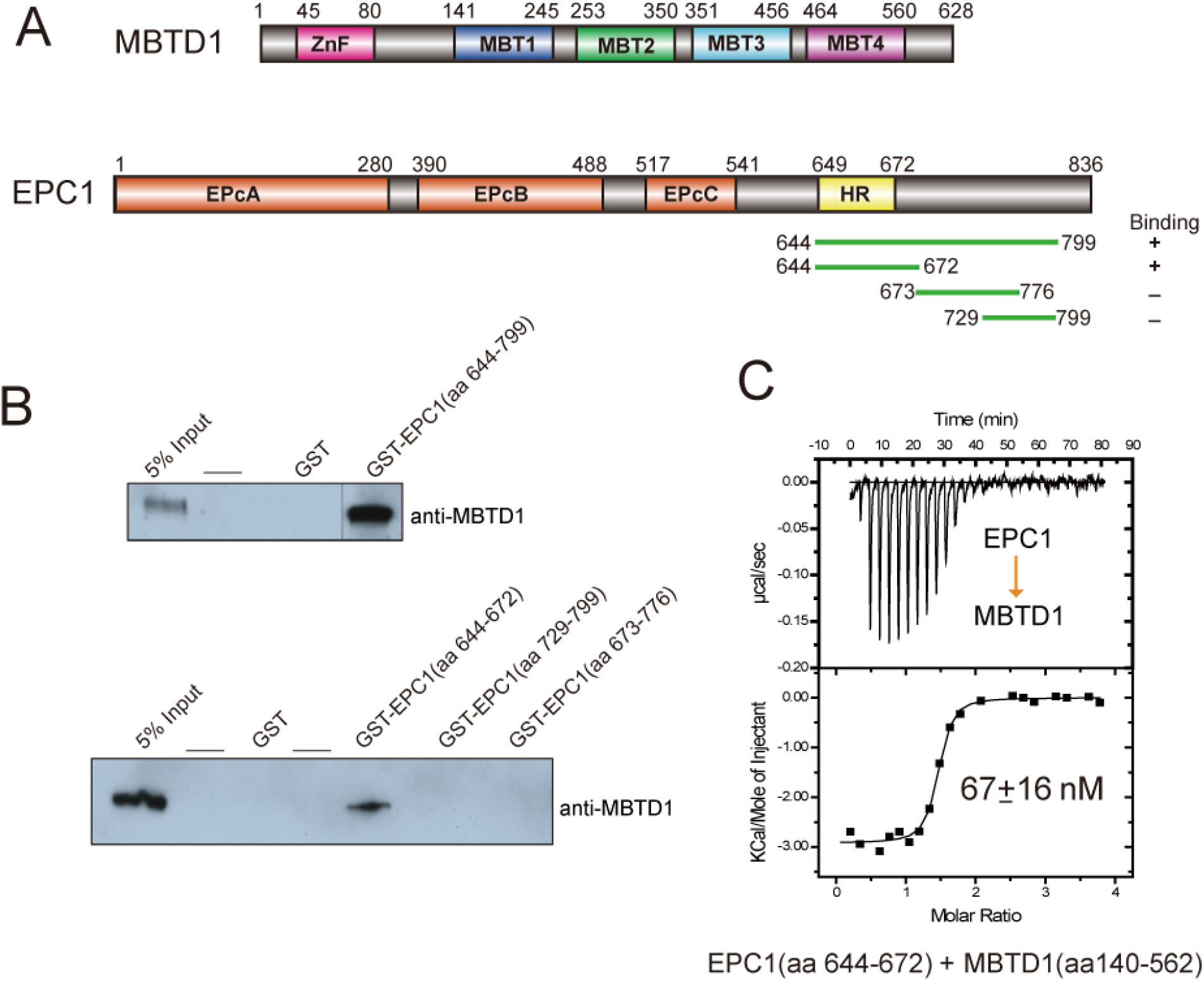
Characterization of the MBTD1-EPC1 interaction. (A) Schematic domain organization of MBTD1 and EPC1. EPc, Epl1 enhancer of polycomb; HR, hydrophobic region; MBT, malignant brain tumor; ZnF, zinc finger. (right bottom) The truncated EPC1 proteins (green lines) used for GST pull-down are labeled with residue numbers. (B) Western blot analysis of GST pull-down using His-MBTD1 incubated with the indicated GST-tagged EPC1 fragments. The amounts of GST-tagged EPC1 proteins used were normalized by Coomassie staining (Fig. S1). (C) The binding of EPC1 to MBTD1 was measured with isothermal titration calorimetry (ITC). The N value is ∼ 1.3. The GST-EPC1 protein was titrated against the MBTD1 protein.

See also Figure S1.

### Overall structure of the MBTD1-EPC1 complex

In order to elucidate how the NuA4/TIP60 acetyltransferase complex engages MBTD1 at the atomic level, we determined the crystal structure of MBTD1_140-562_ in complex with EPC1_644-672_ (**Table S1**). The well-defined electron density allowed us to build residues Lys649-Thr665 of EPC1 and all four MBT domains (MBT1-4) of MBTD1 (**Figure 2A**). Intriguingly, EPC1 folds into a helical structure that locks into a hydrophobic binding groove formed by the MBT1 and MBT2 repeats of MBTD1 (**Figure 2B**), indicating that hydrophobic intermolecular interactions dictate the recruitment of MBTD1 into the NuA4/TIP60 acetyltransferase complex. Specifically, two helices from MBT1 and MBT2 form a V-shaped groove (hereafter named: helical groove) to anchor EPC1. The overall structure of MBTD1 in the complex is highly similar to the previously determined apo structure (PDB: 3FEO) with an RMSD value of approximately 0.36 Å (**Figure S2A**) (Eryilmaz et al., 2009). However, subtle structural divergences are observed in the hydrophobic helical groove of MBTD1 between the apo and MBTD1-EPC1 complex structures. The binding of EPC1 triggers a mild yet significant displacement of the hydrophobic helical groove of MBTD1 compared with the apo structure, especially residues I229-V243, resulting in a relatively more open hydrophobic binding groove upon EPC1 lock-in (**Figure S2B**). Notably, the hydrophobic helical groove in the apo structure of MBTD1 has a high B-factor value, whereas it is well ordered in the complex structure, indicating that the EPC1-binding event can stabilize the hydrophobic helical groove of MBTD1 (**Figure S2C**). Given the facts that 1. only the MBT4 repeat of MBTD1 contains the semi-aromatic cage to recognize H4K20me (Eryilmaz et al., 2009), 2. EPC1-binding does not involve the MBT4 repeat, 3. this binding event does not induce any significant conformational change on the MBT4 repeat structure, it is unlikely that the MBTD1-EPC1 interaction would affect MBTD1’s binding ability towards H4K20me (**Figure 2A**). Consistently, our biophysical analysis demonstrated that MBTD1 displayed similar binding affinities towards H4K20me1 in the presence or absence of EPC1 (**Figure 2C**). Collectively, the above data suggest that the association of MBTD1 with the NuA4/TIP60 core complex does not impair its H4K20me binding ability, thereby conferring this histone mark binding capability to the NuA4/TIP60 acetyltransferase complex and influencing transcription and the DNA repair pathway choice.

**Figure 2.**
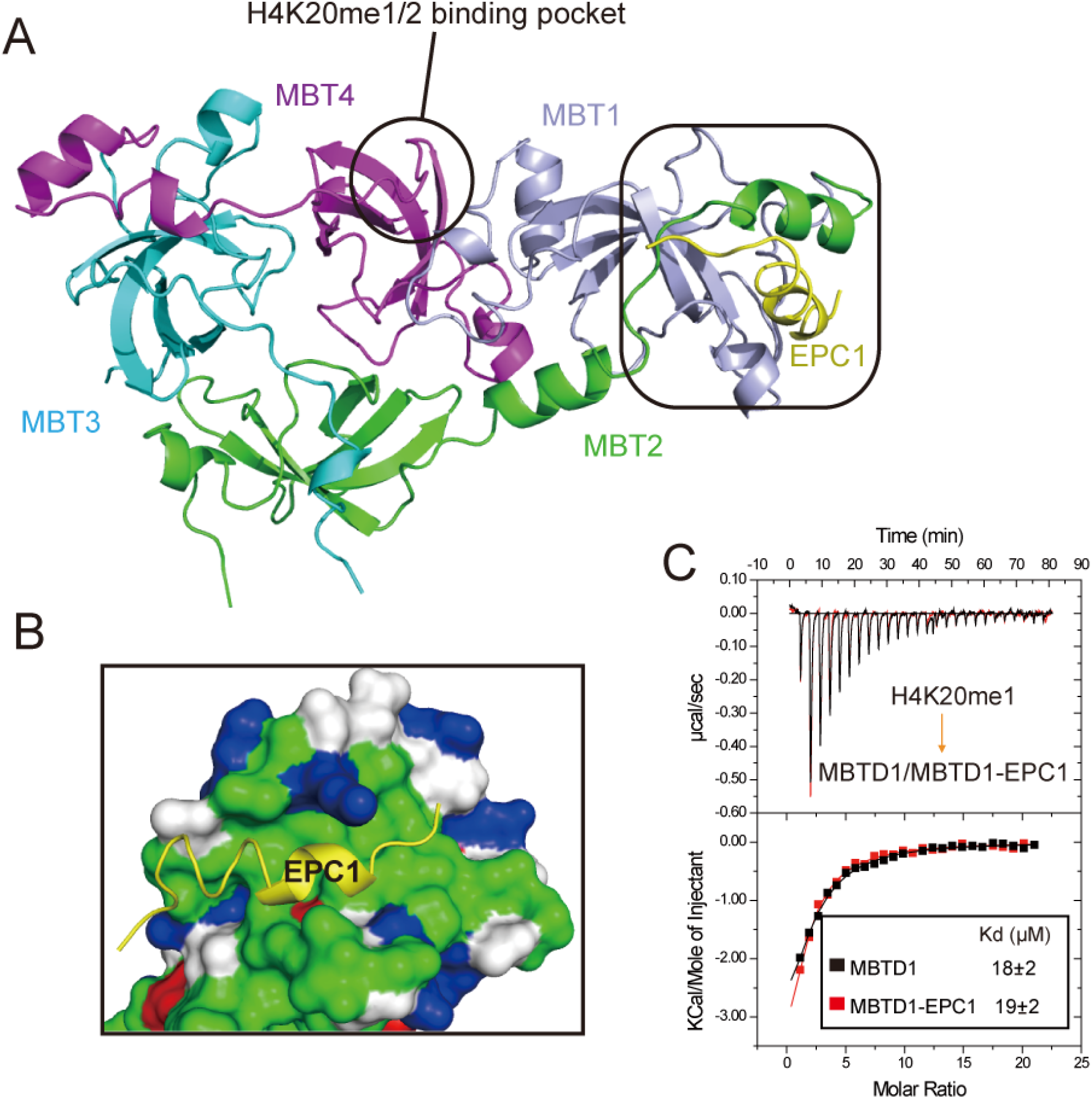
Crystal structure of EPC1 bound to MBTD1. (A) Overall structure of MBTD1 in complex with EPC1. The color scheme is the same as in Fig.1A. (B) Surface representation of MBTD1 showing the hydrophobic pockets for binding of EPC1. Nonpolar residues of MBTD1 are colored green, polar residues grey, positively charged residues blue and negatively residues red. (C) ITC titration curves of H4K20me1 (sequence: AKRHRKmeVLRDN) to MBTD1 (black curve) or the MBTD1-EPC1 complex (red curve). See also Figure S2 and Table S1.

### Recognition mechanism of EPC1 by MBTD1

In the complex structure, EPC1 exhibits an extended conformation and is sandwiched between two helices of MBTD1 (**Figure 2B**). Two hydrophobic leucine residues (Leu651 and Leu663) of EPC1 fit snugly into two separate hydrophobic pockets in MBTD1, respectively (**Figure 3A**). Specifically, Leu663 located at the C-terminus of EPC1 occupies the hydrophobic cleft formed by Trp193, Pro244, Trp255 and Leu259 of MBTD1 (**Figure 3A**). Leu651 from the N-terminus of EPC1 forms hydrophobic interactions with Val232, Leu263 and Ala266 of MBTD1. Similarly, at the center of the binding interface, Ala660 is accommodated by hydrophobic patch created by Leu259, Val260 and Leu263 of MBTD1. Ala656, Phe658 and Ala662 of EPC1 also made hydrophobic interactions with their respective neighboring residues of MBTD1 (**Figure 3A**). In addition to the hydrophobic contacts, hydrogen bonds mediate the intermolecular interactions, further strengthening their association. In particular, the main chain of Leu651 from the N-terminus of EPC1 engages in hydrogen bonds with the main chains of Gly265 and Ala266. At the C-terminus of EPC1, Ala662 and Val664 establish hydrogen bonds with Leu242. Moreover, Ala659 in the middle of EPC1 forms water-mediated hydrogen bonds with Glu179 and Ala236. (**Figure S2F**). Intriguingly, these residues involved in the binding of EPC1 with MBTD1 are largely conserved across different species (**Figures 3B and 3C**), suggesting an evolutionarily conserved recognition mechanism of EPC1 by MBTD1. Taken together, the combined hydrophobic and polar intermolecular interactions are the major driving forces to enlist MBTD1 via EPC1 into the NuA4/TIP60 core complex.

**Figure 3.**
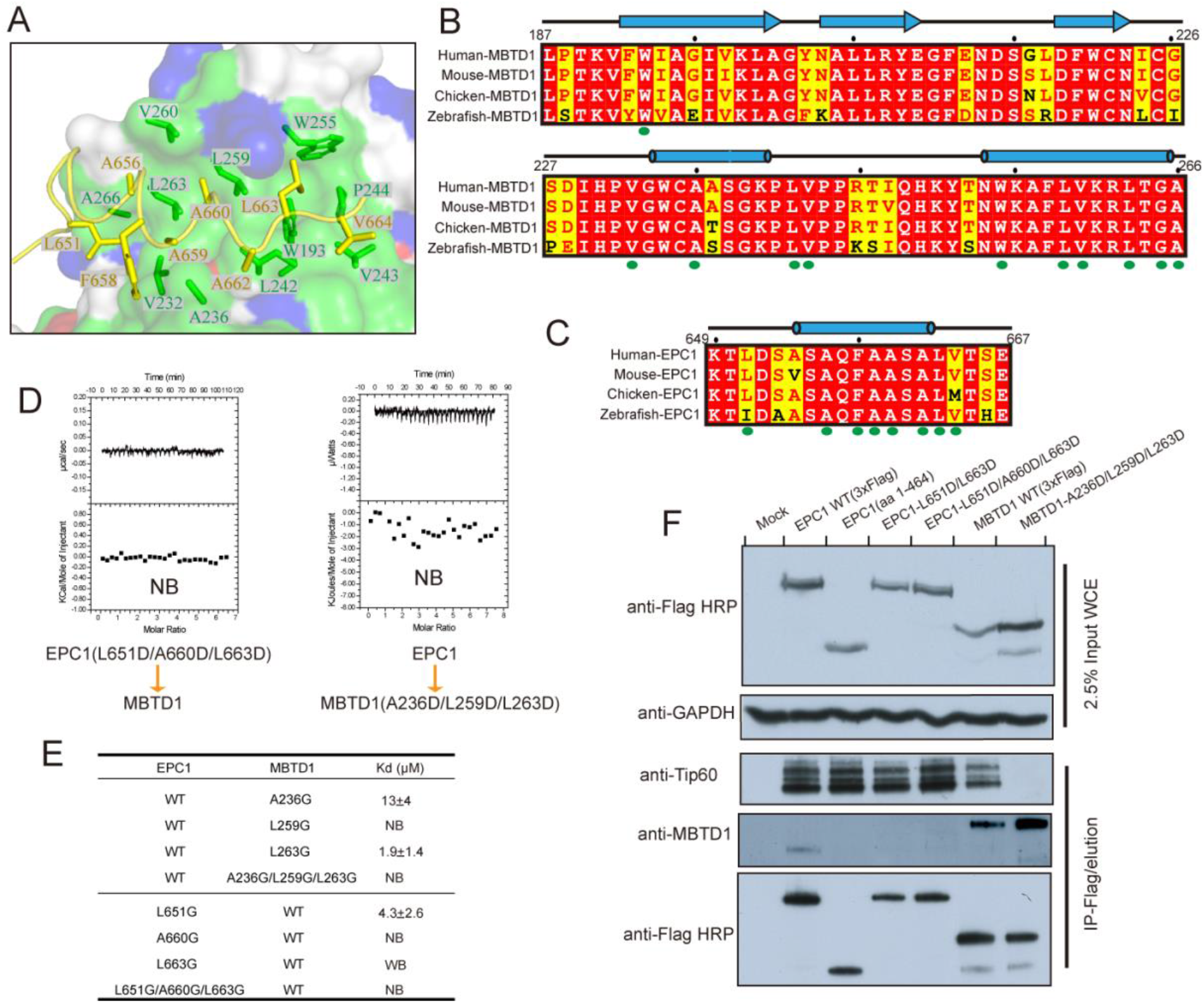
Recognition mechanism of EPC1 by MBTD1. (A) Detailed interactions of MBTD1 with EPC1. The residues involved in the interactions are shown in stick representations, with the same color scheme as in Fig. 2B. (B) Sequence alignment of the helical groove regions of four MBTD1 orthologs. Green circles indicate the residues involved in the MBTD1-EPC1 interactions. (C) Sequence alignment of the MBTD1-binding region of five EPC1 orthologs. (D) Thermodynamic analysis of the interaction between EPC1 triple mutation and MBTD1 (left), and EPC1 and MBTD1 triple mutation (right). NB: No detectable binding. (E) Quantification of the binding affinity between indicated EPC1 and MBTD1 by ITC. NB: No detectable binding. WB: weak binding. (F) K562 isogenic cell lines stably expressing 3xFlag tagged-EPC1 or 3xFlag tagged-MBTD1 with the indicated mutations from the AAVS1 locus. Complexes were immunoprecipitated with anti-Flag antibody, eluted with Flag peptides and analyzed by western blotting with the indicated antibodies. See also Figures S2 and S3.

### Structural comparison of the MBTD1-EPC1 and MBTD1-YY1 complexes

Given that the transcription factor YY1 has been previously demonstrated to directly bind to MBTD1 *in vitro* (Alfieri et al., 2013), we sought to compare the recognition mechanisms of YY1 and EPC1 by MBTD1. Interestingly, MBTD1 employs the same helical groove to capture both EPC1 and YY1, suggesting a conserved scaffold role of the helical groove in the assembly of MBTD1-associated protein complexes (**Figure S2D)**. However, substantial structural differences are observed between MBTD1-EPC1 and MBTD1-YY1. EPC1 folds into a short helix in the MBTD1-EPC1 complex, whereas YY1 adopts a two-stranded β-sheet in the MBTD1-YY1 complex (**Figure S2D**). Furthermore, both the helical groove and YY1 have much higher B-factor values compared with those in MBTD1-EPC1 complex (**Figure S2C**), indicating the flexibility of YY1. Consistently, a much weaker binding affinity has been reported between YY1 and MBTD1 in the micro-molar affinity range, and no detectable binding is observed between YY1 and MBTD1 *in vivo* (Alfieri et al., 2013; Jacquet et al., 2016). Although MBTD1 binds both EPC1 and YY1 predominantly through hydrophobic interactions, several key EPC1-binding residues of MBTD1 (Trp193, Ala236, Val243 and Trp255) are not involved in YY1 binding (**Figure S2E**). Point mutations of these residues significantly impaired the MBTD1-EPC1 association (**Figures 3E and S3**). Hence, the absence of these hydrophobic interactions in the MBTD1-YY1 might explain why MBTD1 displays a much weaker binding to YY1.

### Mutational analyses identified some key residues involved in the MBTD1-EPC1 interaction

To validate the importance of the residues involved in the MBTD1-EPC1 interaction, we introduced point mutations of some key residues into MBTD1 and EPC1 and tested their binding affinities by ITC. Our structural analysis shows that Leu651, Ala660 and Leu663 of EPC1 are involved in hydrophobic interactions with MBTD1, suggesting an important role of these residues in the complex formation. Consistently, the triple mutant EPC1_L651D/A660/L663D_ abolished its binding to MBTD1 (**Figure 3D**). Similarly, the aspartic acid substitutions of the key residues of MBTD1 (A236D/L259D/L263D), identified from the MBTD1-EPC1 complex structure, impaired the MBTD1-EPC1 association (**Figure 3D**). Furthermore, GST pull-down assays showed that the aspartic acid substitutions of some other key residues in the MBTD1-EPC1 interface also significantly dampened the complex formation (**Figures S3A and S3B**). To further discern the residues critical for MBTD1-EPC1 association, we generated glycine substitutions for both EPC1 and MBTD1 proteins. Our ITC data showed that single glycine substitutions of hydrophobic residues significantly reduced or disrupted the MBTD1-EPC1 binding (**Figures 3E and S3C**). Notably, triple glycine mutations abolished the interactions between EPC1 and MBTD1, which is consistent with the aspartic acid substitutions. Collectively, our mutagenesis analysis confirmed the importance of the key residues involved in the hydrophobic interactions between MBTD1 and EPC1 in leashing MBTD1 to the NuA4/TIP60 complex.

To further examine the impacts of the key EPC1 mutations on their abilities to associate with MBTD1 *in vivo*, we generated isogenic K562 cell lines with a single copy of the epitope-tagged EPC1 gene inserted at the *AAVS1* safe harbor locus as described previously (Jacquet et al., 2016). As expected, a C-terminal deletion mutant of EPC1 (EPC1_1-464_) completely lost its ability to associate with MBTD1 in cells (**Figure 3F**). In accordance with the above ITC results, the triple mutant EPC1_L651D/A660D/L663D_ showed complete loss of co-purification with MBTD1, but retained association with Tip60 *in vivo* (**Figure 3F**). Similar experiments with K562 cell lines expressing epitope-tagged MBTD1 confirmed that residues Ala236, Leu259 and Leu263 of MBTD1 are fundamental to MBTD1’s ability to associate with the NuA4/TIP60 complex *in vivo* (**Figure 3F**). Thus, the above results validated the physiological significance of these key residues of MBTD1 and EPC1 for their association in human cell lines.

### Functional implications of uncoupling MBTD1 from the NuA4/TIP60 acetyltransferase complex

MBTD1 has been found to co-localize with other NuA4/TIP60 complex subunits at the transcription start sites of active genes, affecting their recruitment and thereby regulating gene transcription (Jacquet et al., 2016). We next examined the functional significance of the interaction between MBTD1 and EPC1 on gene transcription. The MBTD1 or EPC1 mutants that impair the complex formation exhibited significantly reduced binding to the *RPSA* gene promoter, a *bone fide* target of the NuA4/TIP60 complex, as measured by chromatin immunoprecipitation (ChIP) assay (**Figure 4A**). While a portion of EPC1 is retained, the MBTD1 recruitment is completely lost, in agreement with its dissociation from the complex. Furthermore, MBTD1 and EPC1 have been suggested to function as putative tumor suppressors (Jacquet et al., 2016). Clonogenic survival analysis revealed that transfected cells expressing EPC1 or MBTD1 mutants displayed fewer colony growth compared to the empty vector control (**Figures 4B**, **S4A and S4B**). Notably, significantly more colonies were recovered with the MBTD1 mutant versus its WT version, indicating that MBTD1-dependent growth suppression requires interaction with the NuA4/TIP60 complex at least in part. Collectively, these results confirm the physiological significance of the key residues identified in the MBTD1-EPC1 complex structure.

**Figure 4.**
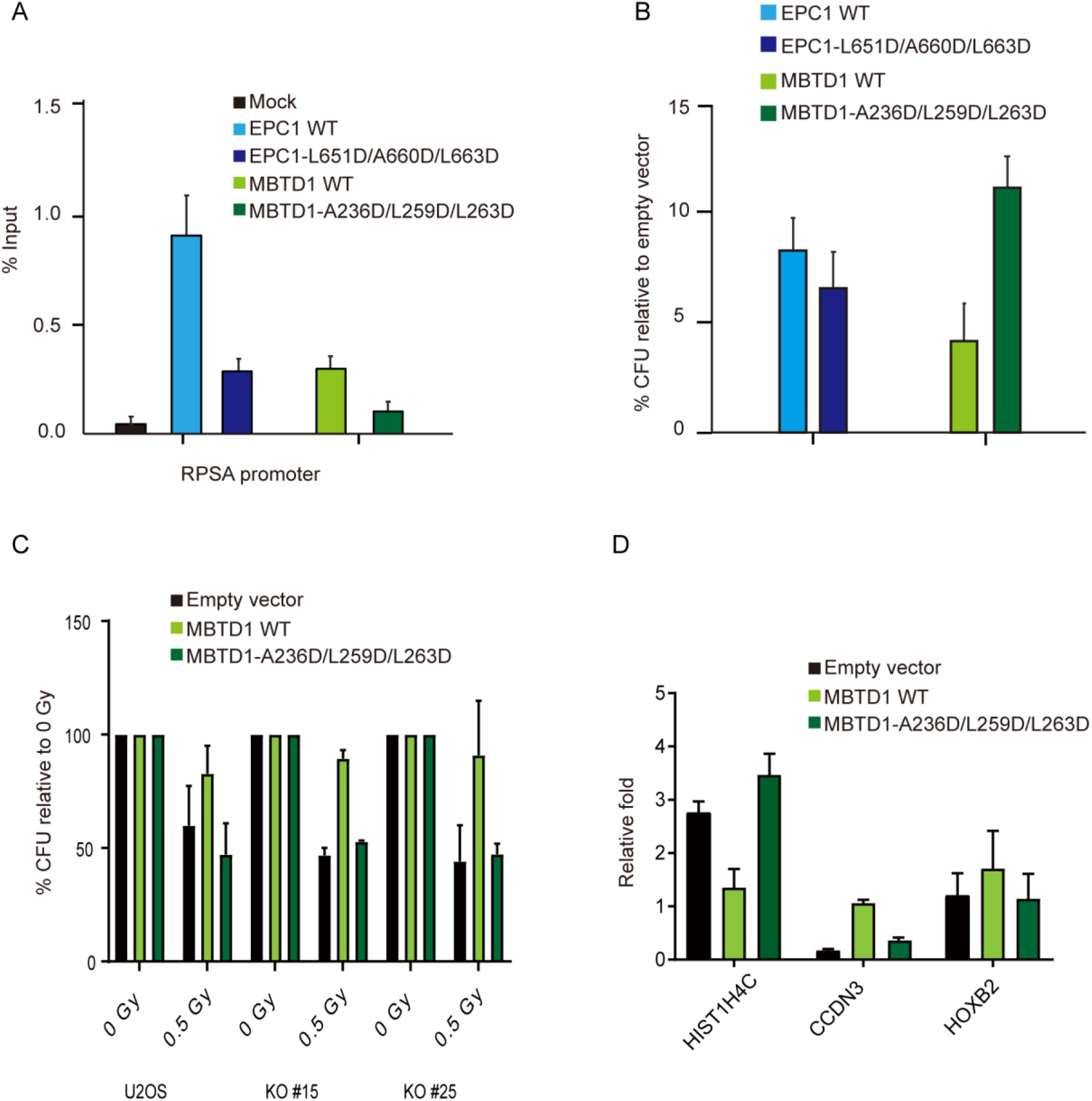
Interaction of MBTD1 and EPC1 impacts gene regulation, DNA damage response and cell growth. (A) MBTD1 or EPC1 mutants affect recruitment to a target gene *in vivo*. Chromatin immunoprecipitations was performed using K562 cells stably expressing MBTD1 or EPC1 mutant proteins. Error bars represent the range of two independent experiments. (B) Clonogenic assay using U2OS cells transfected with the indicated expression vectors or empty vectors. (C) WT MBTD1 but not the EPC1-binding mutant suppresses the DNA damage sensitivity of MBTD1 KO cells. Clonogenic survival assay showing the percentages of viable colonies for each condition post a single dose of gamma-irradiation (0.5Gy) relative to their respective untreated controls. (D) WT MBTD1 but not the EPC1-binding mutant complements the gene regulation defects in MBTD1 KO cells. Expression levels of genes that are dysregulated in MBTD1 KO cells. (Jacquet et al. 2016) as measured by RT-qPCR 48h post-transfection after transfection with empty vector, MBTD1-WT or MBTD1-A236D/L259D/L263D. WT and mutant MBTD1 are expressed to similar levels in this experiment (Fig. S4D). See also Figure S4.

The MBTD1-associated NuA4/TIP60 complex recognizes the H4K20me mark at DSB sites and catalyzes the acetylation of the histone H4 tail and H2AK15 surrounding the sites of broken chromatins (Tang et al., 2013; Jacquet et al., 2016). These DNA damage-signaling events act in concert to favor downstream recruitment of the HR-mediated repair machinery as the choice of DSB repair pathways, blocking at multiple levels the recruitment and retention of the key NHEJ factor 53BP1 to the DSB sites (Jacquet et al., 2016). Given that we have identified the key residues of MBTD1 responsible for its enlistment into the NuA4/TIP60 complex, we aimed to further validate these key residues’ functional importance in mammalian cells. It was shown that overexpression of WT MBTD1 led to mild increase of H2AK15ac signal in MBTD1 knockout cells, whereas overexpression of the MBTD1 mutant failed to produce this effect (**Figure S4C**). Furthermore, post-gamma irradiation clonogenic survival analysis demonstrated that complementation of MBTD1 knockout cells (2 different clones (Jacquet et al. 2016)) by transfection of a WT MBTD1 expression vector rescues the radiosensitivity of MBTD1 knockout cells (**Figure 4C**). In contrast, complementation of the MBTD1 knockout cells with the MBTD1 mutant failed to rescue the radiosensitivity of MBTD1 knockout cells (**Figure 4C**). Interestingly, ectopic expression of MBTD1 seems to also protect WT cells (containing endogenous MBTD1 protein), while the mutant again fails to do so. These results highlight the functional significance of the MBTD1-EPC1 interactions in repairing damaged chromatin in mammalian cells.

Since MBT1 is important for proper expression of specific genes, directly bound by the NuA4/TIP60 complex or indirectly, we next investigated whether WT MBTD1 but not the mutant lacking the EPC1 binding ability rescues the gene regulation defects in MBTD1 knockout cells. The expression levels of *HIST1H4C* and *CCDN3* have been shown to be dysregulated in opposite manners in the MBTD1 knockout cells, *HIST1H4C* being overexpressed while *CCDN3* being underexpressed (Jacquet et al., 2016). We measured the expression levels of these two genes in the MBTD1 knockout cells transfected with empty vector, WT MBTD1 or mutant vectors by RT-qPCR. The *HOXB2* expression is measured as a negative control as it is not affected by the loss of MBTD1. Complementation of the MBTD1 knockout cells with a WT MBTD1 vector rescued their transcription defects, significantly decreasing the expression of *HIST1H4C* while increasing the expression of *CCDN3* (**Figure 4D**). In contrast, the MBTD1 mutant failed to rescue the misregulation of *HIST1H4C* and *CCDN3*, while being expressed at similar levels compared to the WT vector (**Figures 4D** and **S4D**). Thus, the above findings validated the key role of the MBTD1-EPC1 interactions in NuA4/TIP60 acetyltransferase complex-mediated transcriptional regulation.

In brief, our study has elucidated the molecular mechanism by which MBTD1 is enrolled into the dynamic NuA4/TIP60 acetyltransferase complex to regulate transcription and DNA repair pathway choice. These findings will provide valuable information for the development of inhibitors for selectively dissociating MBTD1 from the NuA4/TIP60 acetyltransferase complex, which can have important clinical implications. Drug-induced disruption of the MBTD1 interaction with NuA4/TIP60 could be used to sensitize cancer cells to radiotherapy, alone or in combination with PARP inhibitors. This approach may be better than directly inhibiting Tip60 HAT activity which can lead to much larger side effects on healthy cells.

## Acknowledgements

We are grateful to Amelie Fradet-Turcotte for radiosensitivity clonogenic assays. Diffraction data were collected at the Advanced Photon Source of the Argonne National Laboratory. The SGC is a registered charity (number 1097737) that receives funds from AbbVie, Bayer Pharma AG, Boehringer Ingelheim, Canada Foundation for Innovation, Eshelman Institute for Innovation, Genome Canada through Ontario Genomics Institute [OGI-055], Innovative Medicines Initiative (EU/EFPIA) [ULTRA-DD grant no. 115766], Janssen, Merck KGaA, Darmstadt, Germany, MSD, Novartis Pharma AG, Ontario Ministry of Research, Innovation and Science (MRIS), Pfizer, São Paulo Research Foundation-FAPESP, Takeda, and Wellcome. This work was also supported by the Canadian Institutes of Health Research (CIHR) (FDN-143314 to J.C.) and the Natural Sciences and Engineering Research Council of Canada (NSERC), funding reference number RGPIN-2016-06300 to J.M. J.C. holds the Canada Research Chair in Chromatin Biology and Molecular Epigenetics.

## AUTHOR CONTRIBUTIONS

H.Z., M.D., J.C. and J.M. designed the research. H.Z., M.D., X.S., G.A., H.C., W.T., C.D. and P.L. performed experiments. H.Z., M.D., L.L., J.C. and J.M. wrote the manuscript.

## STAR METHODS

### LEAD CONTACT AND MATERIALS AVAILABILITY

Further contact information and requests for resources and reagents should be directed to and will be fulfilled by the Lead Contact, Jinrong Min. This study did not generate new unique reagents.

### EXPERIMENTAL MODEL AND SUBJECT DETAILS

The plasmid DNAs were amplified in *E. coli* DH5α cells. MBTD1 and EPC1 proteins were expressed in *E. coli* BL21-CodonPlus (DE3)-RIL cells cultured at 37 °C or 18 °C in TB medium. K562 and U2OS cells were obtained from the ATCC and maintained at 37 °C under 5% CO2. K562 were cultured in RPMI medium supplemented with 10% newborn calf serum and GlutaMAX. U2OS cells were cultured in DMEM medium supplemented with 10% fetal bovine serum.

### METHOD DETAILS

#### Protein expression and purification

The MBTD1_140–562_-(GSA)6-EPC1_644–672_ fusion protein construct was cloned into the pET28-MHL vector. Recombinant proteins were expressed in *E. coli* strain BL21-CodonPlus (DE3)-RIL in TB medium overnight at 16 °C. Cells were collected by centrifugation at 4°C and resuspended in 25 mM Tris–HCl pH 8.0, 500 mM NaCl, 5 mM β-mercaptoethanol, 1 mM PMSF. Subsequently, cells were lysed by sonication and debris was removed by centrifugation at 4 °C. The supernatants were loaded onto a Ni-NTA resin (QIAGEN) gravity column and eluted with buffer containing 25 mM Tris-HCl pH 8.0, 500 mM NaCl, 250 mM imidazole, 5 mM β-mercaptoethanol. The proteins were further purified by anion-exchange chromatography and subsequent gel-filtration chromatography in a buffer containing 25 mM Tris-HCl pH 8.0, 150 mM NaCl. Peak fractions were concentrated to 16 mg/mL.

#### Crystallization, data collection and structure determination

Co-crystallization trials of purified MBTD1_140–562_ and EPC1_644-672_ proteins were not successful. Thus, we generated a fusion construct in which EPC1_644-672_ was C-terminally linked to MBTD1_140–562_ using a (GSA)_6_ linker. Remarkably, the MBTD1-EPC1 fusion protein yielded high-quality crystals that diffracted to a resolution of 1.9 Å and enabled us to solve the complex structure. Crystal screening was performed by sitting-drop vapor-diffusion at 18 °C. Crystals of MBTD1-EPC1 were obtained using a reservoir solution composed of 0.1 M sodium citrate tribasic dihydrate pH 5.0, 10% w/v polyethylene glycol 6,000. The crystals were soaked in a cryoprotectant of crystallization solution supplemented with 20% glycerol before being flash-frozen. Diffraction data were collected at beam line 19-ID of the Advanced Photon Source and reduced with XDS (Kabsch, 2010) and AIMLESS (Evans and Murshudov, 2013). The structure was solved by molecular replacement with coordinates from PDB entry 3FEO (Eryilmaz et al., 2009). The model was iteratively refined with REFMAC (Murshudov et al., 2011) and rebuilt with COOT (Emsley et al., 2010). Model geometry was evaluated with MOLPROBITY (Chen et al., 2010).

#### GST pull-down

EPC1 fragments with the indicated boundaries were cloned into the pET28GST-LIC vector that encodes an N-terminal GST-tag. All point mutations were generated using the Quick Change Site–directed mutagenesis system (Stratagene, La Jolla, CA). All GST-tagged proteins were expressed in BL21(DE3) cells and purified by GST affinity chromatography (Glutathione agarose). Proteins were then dialyzed overnight against dialysis buffer (25 mM Tris-HCl, pH 8.0, 150 mM NaCl, 1 mM β-mercaptoethanol). The GST-tagged proteins were first incubated with glutathione resin for 1 hour at 4 °C in dialysis buffer. His-tagged MBTD1 proteins were incubated with the beads for 30 min at 4 °C. The samples were washed four times with 1 mL dialysis buffer to remove excess unbound protein, and then analyzed through Coomassie staining of SDS-PAGE or western blotting. Anti-MBTD1 (Abcam Ab116361) antibodies were used for western blotting at 1:1000 dilution.

#### Isothermal titration calorimetry

Proteins were expressed and purified as described above. The ITC experiments were performed on a Microcal VP-ITC (Microcal, Amherst, MA) at 25 °C. All measurements were carried out at a stirring speed of 307 rpm, in 25 mM Tris-HCl, pH 8.0, 150 mM NaCl. For H4K20me1 binding assay, the H4K20me1 peptide (1 mM) was titrated into the MBTD1 or MBTD1-EPC1 fusion protein solution (10 μM) with 26 injections. For the MBTD1-EPC1 binding assay, each titration was carried out by injecting the EPC1 WT or mutants proteins (0.2 mM) into the cell containing MBTD1 WT or mutants proteins (10 μM). The ITC experiments were repeated at least two times. The binding data were subsequently analyzed using Origin (MicroCal).

#### Generation of isogenic cell lines and affinity purification of complexes

K562 cells were used to express various 3xFlag-tagged EPC1 proteins including EPC1_WT_, EPC1_1-464_, EPC1_L651D/L663D_, and EPC1_L651d/A660d/L663d_ as well as 3xFlag-tagged MBTD1_WT_ and MBTD1_A236d/L259d/L263d_ from the AAVS1 safe harbor using ZFN-mediated insertion as described previously (Dalvai et al., 2015). After whole cell extraction, anti-Flag immunoprecipitation was performed with anti-FLAG agarose affinity gel (Sigma M2), followed by elution with 3xFLAG peptide (200 μg/ml from Sigma in the following buffer: 20 mM HEPES pH7.9, 100 mM KCl, 0.1% Triton X-100, 20% glycerol, 100 μM ZnCl2, 1 mM DTT and supplemented with proteases, deacetylases and phosphatases inhibitors).

Flag eluates from the above preparations were normalized following SDS-PAGE and transfer onto a nitrocellulose membrane. The following antibodies were used at the indicated dilution: anti-FLAG-HRP conjugate (Sigma M2, 1:5000), anti-Tip60 (Abcam 137518, 1:1000); anti-MBTD1 (Abcam ab116361, 1:1000); anti-GAPDH (ThermoFisher, 39-8600, 1:10000).

#### ChIP-qPCR Assay

For anti-FLAG ChIP, 1 mg of cross-linked chromatin from K562 cells was incubated with 10 μg of anti-FLAG antibody (Sigma, M2) or anti-IgG antibody (Millipore-PP64) pre-bound on 300 μl of Dynabeads Prot-G (Invitrogen) overnight at 4 °C. The beads were washed extensively and eluted in 0.1% SDS, 0.1 M NaHCO3. Crosslink was reversed with 0.2 M NaCl and incubation overnight at 65 °C. Samples were treated with RNase and Proteinase K for 2 h and recovered by phenol chloroform extraction and ethanol precipitation. Quantitative real-time PCRs were performed on a LightCycler 480 (Roche) with SYBR Green I (Roche) to confirm specific enrichment at the RPSA locus. Ratios of immunoprecipitated signal versus input are based on duplicate experiments and errors bars represent the range. Primers used: AGAAAGCGGGCTAACATCCT; GCTGTGCTGTCACCACTTGT.

#### Clonogenic growth assay

EPC1_WT_, EPC1_L651D/A660D/L663D_, MBTD1_WT_, MBTD1_A236D/L259D/L263D_ or empty vectors were transfected into U2OS cells which were then seeded onto dishes in duplicates. Subsequently, puromycin [1μg/μl] was added to the culture media at 48hours post-transfection for selection. After ten days of culture, the colonies formed on the dishes were fixed with methanol, followed by staining with crystal violet (Sigma C6158) and washing with distilled water. Images of the dishes were then taken.

#### Gamma-irradiation and clonogenic survival assay

MBTD1-3xFlag, MBTD1-A236D/L259D/L263D-3xFlag mutant or an empty vector were transfected into U2OS cells or MBTD1 KO cell lines previously described (Jacquet et al. 2016). Cells were then seeded in 10cm dishes. Puromycin was added to the culture media [1μg/ml] 48hrs post-transfection for two days of selection. One day before irradiation, the cells were counted to plate 500 cells per condition in 6-well plates. Irradiation with 0 or 0.5 gray was performed using a Cellrad Faxitron irradiator. After 12 days of culture, the colonies formed were fixed with methanol, followed by staining with crystal violet (Sigma C6158) and washes with distilled water. Images were then taken, and colonies counted.

Western blot analysis of whole cell extracts from the above transfections were done to verify equivalent expression of WT and mutant proteins. The following antibodies were used at the indicated dilution: anti-FLAG-HRP conjugate (Sigma M2, 1:5000) and anti-GAPDH (ThermoFisher, 39-8600, 1:10000). The error bars represent the range based on independent experiments.

#### Chromatin extracts

Chromatin-enriched extracts were prepared as described before (Gwak et al., 2016). Briefly, U2OS cells were transfected with WT MBTD1-3xFlag, MBTD1-A236D/L259D/L263D-3xFlag mutant or an empty vector (4μg) using Lipofectamine 2000. 2 days post-transfection, puromycin was added [1μg/ml]. 48hrs later cells were harvested, washed once with PBS 1X and resuspended in buffer 1 (50mM Tris-HCl pH7.5, 100mM NaCl, 1mM EDTA, 1mM DTT, 1X protease inhibitor cocktail (Sigma-P8340), 1mM PMSF, 20mM N-ethylmaleimide, 10mM NaB, 10mM NaF). Cell suspension was incubated 5 min on ice, then centrifuged at 1000g for 15 minutes. Supernatant was discarded. Nuclei were then resuspended in buffer 2 (50mM Tris-HCl pH7.5, 300mM NaCl, 1mM EDTA, 5mM CaCl_2_, 1mM DTT, 1X protease inhibitor cocktail, 1mM PMSF), incubated 10 min on ice and sonicated 3×30 seconds. Chromatin enriched extracts were finally clarified by centrifugation at 1000g for 20 min. The following antibodies were used for Western blotting at the indicated dilution: anti-H2AK15ac (Abcam ab101447, 1:1000); anti-H4penta-Acetyl (Upstate 06-946, 1:3000); anti-H3 (Abcam 1791, 1:10 000); anti-H4 (Abcam 7311, 1:5000).

#### Reverse Transcription-qPCR

WT MBTD1-3xFlag, MBTD1-A236D/L259D/L263D-3xFlag mutant or an empty vector were transfected in a MBTD1 KO U2OS cell line (#15 (Jacquet et al. 2016). Puromycin [1μg/ml] was added 48hrs post-transfection and 48hrs later total RNA was extracted with the RNeasy Plus Mini kit (Qiagen). 500 ng of RNA was reverse transcribed by oligo-dT and random priming into cDNA with a qScript cDNA SuperMix kit (QuantaBio-VWR), according to the manufacturer’s instructions.

Quantification of the amount of cDNA was done with SYBR Green I (Roche) on a LightCycler 480 (Roche) for real-time PCR, using primers specific for *HIST1H4C* and *CCDN3* coding regions, two genes that are affected in MBTD1 KO cells (Jacquet et al. 2016). The *HOXB2* gene was used as negative control as it is not affected in MBTD1 KO cells. The *36B4* gene was used as a housekeeping internal control for normalisation. The error bars represent the range based on independent experiments. The oligonucleotide sequences used for expression analysis by RT-qPCR are listed below:

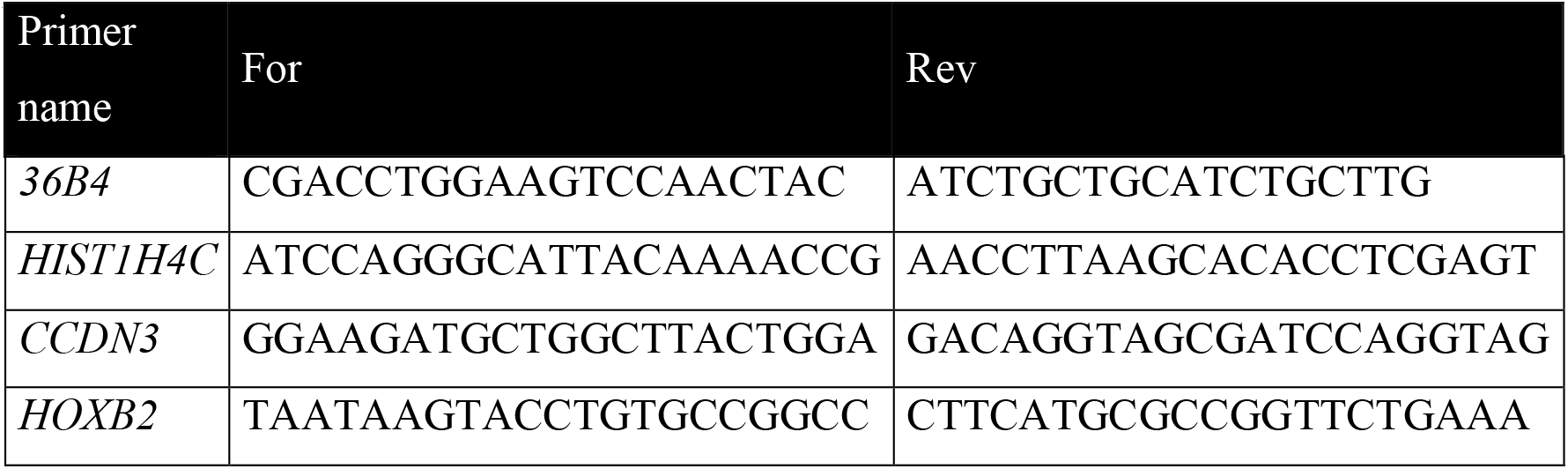

### QUANTIFICATION AND STATISTICAL ANALYSIS

Statistics generated from X-ray crystallography data processing, refinement, and structure validation are displayed in Table S1. ChIP data in Figure 4A are shown as % of input chromatin signal subtract by the IgG background. Clononegic survival assay in Figure 4B and 4C are shown as the mean of the number of colonies relative to the empty vector of two independent experiments. The Figure 4D represents the mean of the quantification of cDNA of specific genes relative to the housekeeping gene 36B4 of two independent experiments. The error bars of all these figures represent the range.

### DATA AND CODE AVAILABILITY

The accession number for the coordinates and structure factors for MBTD1-EPC1 reported in this paper is PDB: 6NFX

